# Collinearity of alpha-helices or beta strands in membrane proteins causes a characteristic peak centred on 4.9 Å resolution in diffraction intensity profiles, inducing higher diffraction anisotropy

**DOI:** 10.1101/2021.12.07.471609

**Authors:** Juliette Martin, Xavier Robert, Patrice Gouet, Pierre Falson, Vincent Chaptal

## Abstract

Diffraction anisotropy is a phenomenon that impacts more specifically membrane proteins, compared to soluble ones, but the reasons for this discrepancy remained unclear. Often, it is referred to a difference in resolution limits between highest and lowest diffraction limits as a signature for anisotropy. We show in this article that there is no simple correlation between anisotropy and difference in resolution limits, with notably a substantial number of structures displaying various anisotropy with no difference in resolution limits. We further investigated diffraction intensity profiles, and observed a peak centred on 4.9Å resolution more predominant in membrane proteins. Since this peak is in the region corresponding to secondary structures, we investigated the influence of secondary structure ratio. We showed that secondary structure content has little influence on this profile, while secondary structure collinearity in membrane proteins correlate with a stronger peak. Finally, we could further show that the presence of this peak is linked to higher diffraction anisotropy.

**Synopsis:** Membrane protein diffraction anisotropy originates from a peak at 4.9 Å resolution in intensity profiles, due to secondary structure collinearity.

## 1. Introduction

Membrane proteins and soluble proteins differ in their 3D arrangement of amino-acids. For their insertion in the membrane, membrane proteins harbor a series of amino-acids displaying significant hydrophobicity towards the outside of the protein, facing the lipid tails of the membrane. These amino-acids fold into typical secondary structures, being either all alpha helical, or all beta strands that form a beta barrel.

For their biochemical and biophysical studies, membrane proteins often need to be extracted from the membrane and stabilized in solution using amphipathic molecules (detergents, amphipols, nanodiscs, polymers, etc.) (Johansen *et al*., 2021, Dufourc, 2021, Krishnarjuna *et al*., 2020, Salnikov *et al*., 2019, Ravula *et al*., 2019, Overduin & Esmaili, 2019, Oluwole *et al*., 2017, Dörr *et al*., 2016, Marconnet *et al*., 2020, Nguyen *et al*., 2018, Matar-Merheb *et al*., 2011, Chae *et al*., 2010, Bayburt & Sligar, 2010). This can have in turn an influence on their studies, for example to crystallize a membrane protein, detergents prevent lateral packing thus decreasing the possible amount of crystal contacts (Robert *et al*., 2017). Alternatively, crystallization can be performed *in-meso*, inducing a different type of crystal packing (Caffrey, 2015, Russo Krauss *et al*., 2013). Surprisingly, membrane proteins diffract X-rays differently than soluble proteins, resulting in higher diffraction anisotropy at a given resolution. In our earlier study, we found that this fact is seen regardless of the nature (type of membrane insertion or function) of the membrane protein constituting the crystal, or the type of crystal packing and arrangement (Type I *in meso, vs* Type II in detergent), but we were unable to identify the root cause for this discrepancy. Only their membranous nature is responsible for this phenomenon as nothing can distinguish them in respect to anisotropy (Robert *et al*., 2017).

Protein crystals diffract X-rays following specific intensity profiles. In 2003, an experimental curve was created from an ensemble of protein crystals diffracting to very high resolution, from protein of various shapes and folds, and thus representing a “typical” protein diffraction profile (Popov & Bourenkov, 2003). On this typical intensity profile, several zones, or resolution regimes, have been assigned with the presence of specific protein characteristics like secondary structures, the presence of nucleic-acids, or ligands (Morris *et al*., 2004, Zwart & Lamzin, 2003). Diffraction anisotropy is defined as a variation from this experimental curve (Read & McCoy, 2016, McCoy *et al*., 2007).

Diffraction anisotropy is also often described as a difference in resolution limits, creating an ellipsoidal type diffraction pattern rather than a circular one (Global_Phasing_Limited, 2018, Chaptal *et al*., 2016, Sawaya, 2014, Chacko *et al*., 2012, Strong *et al*., 2006, Sheriff & Hendrickson, 1987, Evans & Murshudov, 2013). Diffraction anisotropy is measured/calculated on principal axis forming the ellipsoid, and summed in absolute numbers to give an overall value. This difference in resolution limits of a data set has a large impact on data completeness in the high-resolution shells, which affects electron density map quality and results in more noise in difference maps. Diffraction anisotropy also impacts map quality and could create artifacts or deformation of helices (Global_Phasing_Limited, 2018). Often, the two phenomena co-exist but not always, in our experience. To investigate further this relationship, and the link between the two, we downloaded the whole Protein Data Bank (wwPDB_consortium, 2018) for which we calculated anisotropy and difference in resolution limits for each entry, and submitted them to comparison. We found no simple correlation, even though these two events indeed often co-happen. We then analysed intensity profiles of membrane or soluble proteins and identified a peak at 4.9Å resolution, more prevalent for membrane proteins. We could assign this peak to not only the presence of secondary structures, but rather the collinearity of alpha helices and beta strands in membrane proteins. We finally related the presence of this peak to the distribution of diffraction anisotropy. We conclude on the influence of secondary structure elements collinearity in membrane proteins on diffraction anisotropy.

## 2. Methods

### 2.1. PDB data mining and processing

A local copy of the RCSB Protein Data Bank (PDB) (wwPDB_consortium, 2018) was made including all the deposited structures in PDB formatted coordinate files as well as all the crystallographic structure factors in mmCIF format, as of November 6th, 2020. This local copy was processed using software from the CCP4 software suite version 7.0 (Winn et al., 2011, Berman et al., 2000) and in-house automated Linux script in a similar way as described in (Robert et al., 2017). The anisotropic delta-B value, denoted Aniso_B(I) was computed with Phaser while the resolution limits for each of the 3 principal axes of the anisotropic ellipsoid were determined using the program Truncate from CCP4. The ‘DELTA_RES’ value consists in the subtraction of the lowest resolution limit from the highest one. Finally, intensity profiles, divided in 20 resolution bins, were calculated using the raw anisotropy data listed by the ‘ANO’ mode of Phaser.

From this database, we also extracted, or computed when not available, an ensemble of usual data such as resolution, space group, solvent content, etc. We further calculated the content in secondary structure elements (helices, beta strands, turns and random coils) with the program DSSP (Touw *et al*., 2015, Kabsch & Sander, 1983) together with their ratio in the asymmetric unit. The entries were then divided between soluble and membrane proteins, the later based on interrogation of the ‘Membrane proteins of known structures’ database (http://blanco.biomol.uci.edu/mpstruc/) leaded by S.H. White (University of California, Irvine) and the PDBTM database (Kozma *et al*., 2013).

Fully embedded membrane proteins (‘MB_EMB’) are membrane proteins for which no part is significantly protruding on either side of the membrane. This set of proteins have been identified in our previous study (Robert *et al*., 2018, 2017) by manually checking each membrane protein entry and these identified entries have been directly used in the present study.

### 2.2. Database curation

The curation process was to keep all data for which intensities were available (*i*.*e*., all data for which Aniso_B(I) could be computed). Also, in order to compare reasonably well-behaved structures, only data diffracting to less than or equal to 5 Å resolution were kept, and anisotropic delta-B values above 150 Å^2^ were rejected. Thus, our final dataset consisted of 48,149 records divided in two subsets of 47,065 and 1,084 entries (soluble and membrane proteins, respectively).

### 2.3. Statistical analysis

Based on our previous analysis of diffraction anisotropy across the entire PDB, we performed both parametric (Student’s T-test with Welch correction) and non-parametric (Mann-Whitney) statistics on each question where statistics can be applied. In the case of multiple comparison, Analysis of variances (ANOVA) was used to compare variances between groups, and Kruskal-Wallis for analysis of variances on ranks. Caution was used to enforce the same significance statement for parametric and non-parametric statistics before affirming the claim in the manuscript. Statistical analyses and charts rendering were conducted using Excel 2019 (Microsoft Corporation) and PRISM 9 (GraphPad Software, Inc).

Smoothed profile for diffraction intensity profiles. On plots showing intensity profiles as a function of resolution, smoothed profiles were estimated by fitting piecewise cubic polynomials (natural splines) using the splines package in R(R Core Team, 2016) and displayed using ggplot2(Wickham, 2016).

### 2.4. Collinearity determination of secondary structures

The program DSSP (Touw *et al*., 2015, Kabsch & Sander, 1983) was used to map above-mentioned secondary structure elements on each residues of all entries of our dataset. The helix geometry was analyzed using the ‘Helix packing software’(Dalton *et al*., 2003) as follows. We considered the local axes, computed for four consecutive residues in helix, and defined a collinearity index as the percentage of inertia along the first principal component after the principal component analysis of the local axes. The collinearity index varies between 0.3 (random orientation of local axes) and 1 (all axes perfectly colinear).

## 3. Results and Discussion

### 3.1. Very weak correlation between diffraction anisotropy and difference in resolution limits

First, we wanted to evaluate the link between diffraction anisotropy and the difference in resolution limits. To this end, we followed the same procedure as we established in (Robert *et al*., 2017); we downloaded the whole PDB, and crossed it with two databases of membrane protein structures (‘Membrane proteins of known structures’ database (http://blanco.biomol.uci.edu/mpstruc/) and the PDBTM database(Kozma *et al*., 2013)) to sort out soluble and membrane proteins. For each structure, we selected only the one for which structure factors were available in the form of intensities. We then calculated diffraction anisotropy on intensities using Phaser(Read & McCoy, 2016). We also extracted resolution limits along principal reciprocal space directions using the UCLA anisotropy server(Strong *et al*., 2006), identifying the resolution at which the signal/noise ratio was falling below 2, and calculated the difference between the highest and lowest resolution limits. Figure 1 reports diffraction anisotropy (Aniso_B(I)) as a function of difference in resolution limits (DELTA_RES). For all structures (Figure 1A), there is an apparent spread of anisotropy values for each range of difference in resolution limits. There is a trend to more anisotropy for larger difference is resolution limits, which corresponds with the general idea described in the introduction. But there are also many cases that do not follow this behaviour, showing either very low or very high anisotropy for the same value of difference in resolution limits. In addition, there are many cases showing a wide range of anisotropy with no difference in resolution limits (Figure 1B). To avoid pitfalls linked with constraints on cell axis of higher symmetry space groups, we investigated structures belonging to space group P 1 21 1 only (Figure 1C,D). Indeed, this space group in well populated and has minimal constraints on cell parameters. The data in P 1 21 1 scatter similarly to the whole population of structures, and show a substantial number of structures with no difference in resolution limits while having a large spread in anisotropy values. All these observations led us to reconsider the correlation between anisotropy and difference in resolution limits, but rather appreciate these two variables as distinct while often converging.

**Figure 1.**
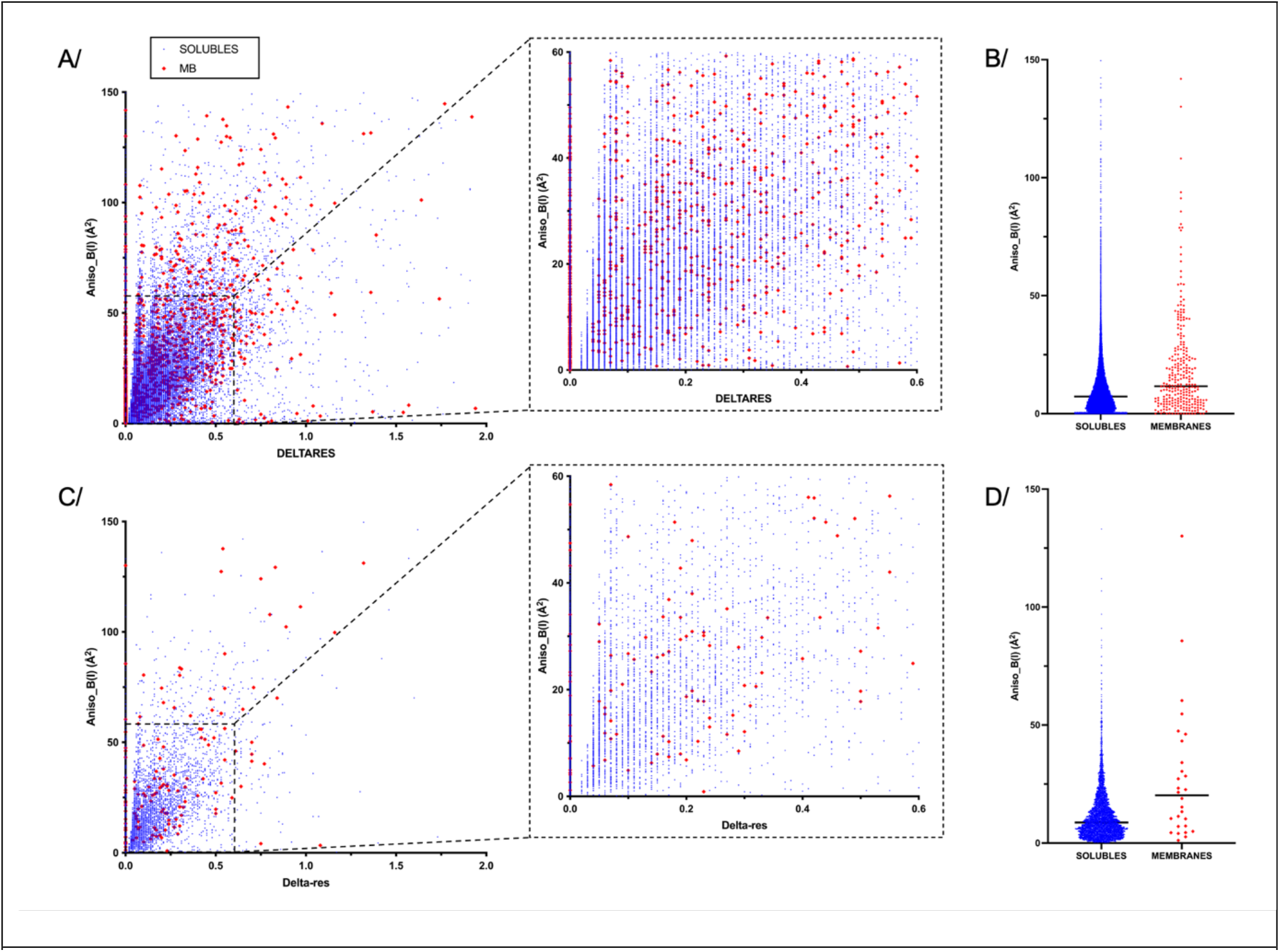
Diffraction anisotropy as a function of difference in resolution limits. A/ 2D plot representing anisotropy values in Å^2^ (calculated on Intensities) as a function of the difference in resolution limits (deltares). The right panel is a zoom on the section between 60Å^2^ Aniso_B(I) and 0.6 deltares constituting the most populated area of the graph. Each point represents a PDB entry; soluble proteins are in blue, membrane proteins are in red. B/ Scatter plot for all entries with no difference in resolution limits. The median is shown as a black bar. C/ and D/ are same as A/ and B/ for entries belonging to space group P 1 21 1.

### 3.2. Intensity profiles for soluble and membrane proteins

At the core, diffraction anisotropy is defined as a deviation from the experimental observation made by Popov & Bourenkov on 72 crystals of soluble proteins diffracting to very high resolution(Popov & Bourenkov, 2003). We thus next investigated diffraction intensity profiles for membrane proteins and compared them with soluble ones (Figure 2). Intensity profiles were calculated for each entry by Phaser, which normalises them to unity compared to the curve described by Popov & Bourenkov. A deviation from unity shows a deviation from the experimental curve. Figure 2 displays the intensity profiles for each entry, overlaid, as a function of resolution. Each line corresponds to a PDB entry. Some variability is seen at low resolution, as expected in this region. Then profiles follow unity with a small degree of variation for soluble proteins, as was also observed previously for a chosen set of soluble proteins (Morris *et al*., 2004). The smoothed profile is highlighted in blue (see M&M section for a description of the fit). Membrane protein profiles show a remarkable difference in the secondary structure elements zone(Morris *et al*., 2004), with a large peak centred on 4.9 Å and two valleys at 3.5 and 6.75 Å deeper that soluble protein profiles. The peak can be more or less pronounced in between proteins, but is always present. Fully embedded membrane proteins show a more pronounced profile. This observation is important and will be discussed more in depth later. One important fact about embedded membrane proteins is that they display a particular fold, nearly all alpha helical or all beta strands, oriented normal to the membrane plane.

**Figure 2.**
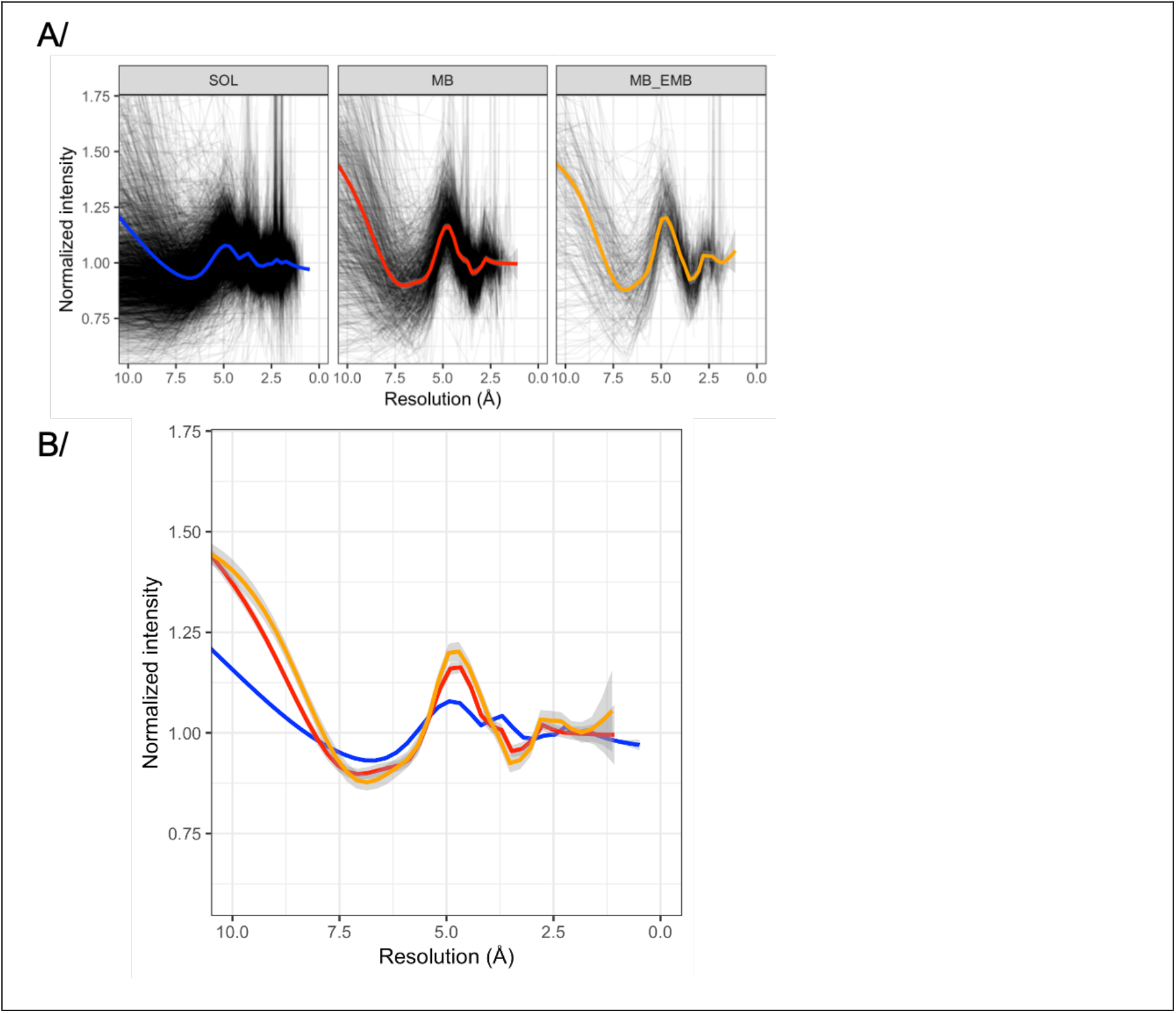
Intensity profiles for soluble and membrane proteins. A/ Intensity profiles normalized to unity (black lines) as displayed as a function of resolution, for soluble proteins (SOL), membrane proteins (MB) and membrane proteins embedded into the membrane without any extra-membranous parts (MB-EMB). For soluble proteins, only 3000 randomly chosen entries were displayed for clarity; each line corresponds to a PDB entry. Blue, red and orange traces correspond to the smoothed intensity profile for all entries. B/ Smoothed intensity profiles superposed for comparison.

### 3.3. Presence of secondary structures alone is not enough to describe the peak centred on 4.9 Å resolution

Secondary structure elements in folded protein structures produce characteristic repeated distances in the 4.5-7 Å range (Morris *et al*., 2004, Zwart & Lamzin, 2003, Morris & Bricogne, 2003). Thus, the more secondary structure elements, the larger the amount of distance repeats in these resolution regimes. We investigated the influence of secondary structures on the formation of the peak centred on 4.9 Å. Soluble and membrane proteins were separated by percentage of secondary structure elements constituting their fold and intensity profiles displayed (Figure 3). Separation on the percentage of alpha helices alone (Figure 3A) reveals that for membrane proteins the peak is very present even for low content in alpha helices. The peak, and the inflections around it, are more pronounced as the proportion of alpha helices increases. For soluble proteins however, the trend reverses with a small peak detectable for low alpha contents that then flattens for higher alpha percentage. A similar trend is observed when the separation is performed on beta strands only for membrane proteins. Soluble proteins show a peak for high beta strand content, albeit smaller in height compared to membrane proteins (Figure 3B).

**Figure 3.**
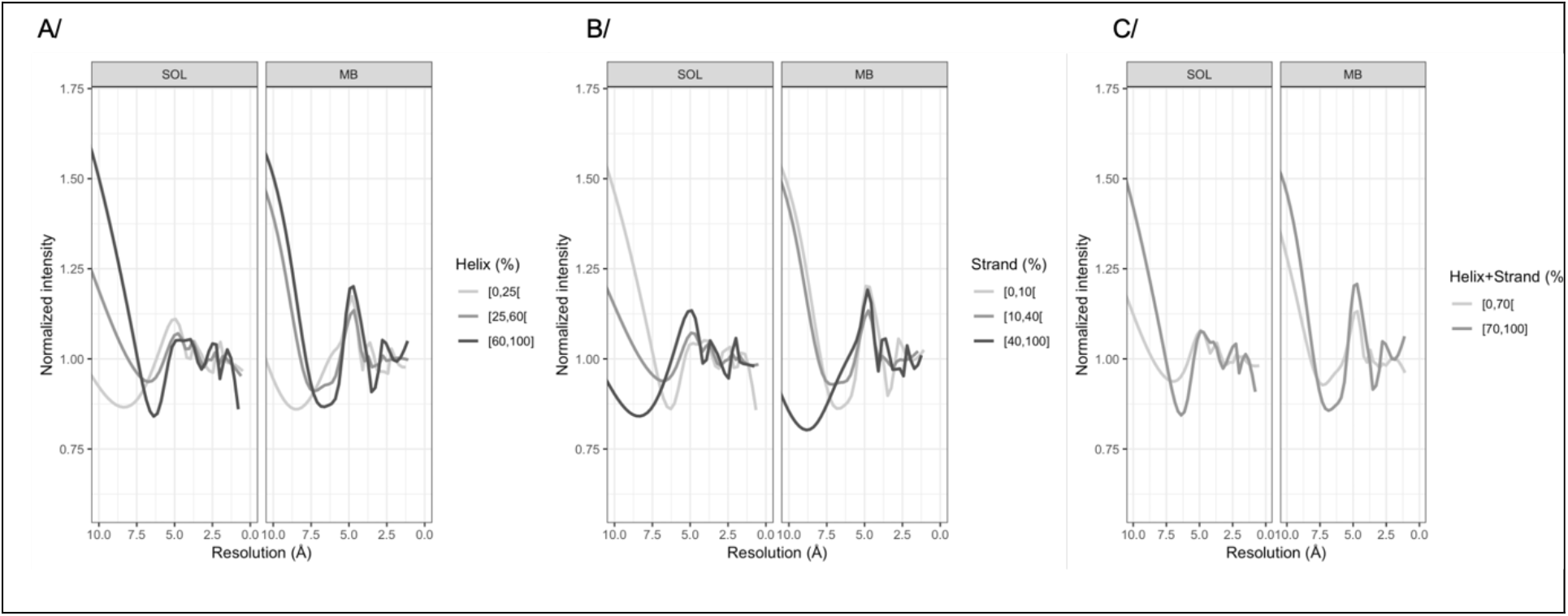
Influence of secondary structure proportion on intensity profiles. A/ separation on helix proportion alone. B/ separation on beta strand proportion alone. C/ separation on the content of helices and strands. Soluble (SOL) and membrane (MB) proteins were separated by secondary structure proportion with smoothed traces overlaid and colored according to secondary structure proportion. Limits were chosen based on secondary structure distribution (not shown).

Since proteins are mostly constituted of a combination of alpha helices and beta strands, and a low content in one element is usually balanced by a higher content of the other, the influence of all secondary structure elements on intensity profiles was investigated (Figure 3C). While on one hand the trend is confirmed for membrane proteins, for soluble proteins on the other hand the influence of both secondary structure elements counter balance and the average intensity profile is close to unity for all proportions of secondary structure content. All these information together show that the ratio of secondary structures *per se* is not the sole source of an increase in intensity profile.

### 3.3. Selected soluble proteins for a peak at 4.9Å resolution show a higher proportion of beta strands

In the intensity profile for soluble proteins a variability was observed (Figure 2A), and a small number of structures were recognized to display the same intensity profile as membrane proteins, with a marked peak centred on 4.9Å resolution. We selected out these soluble proteins on the criteria to have an intensity ratio over 1.25 at 4.9Å resolution (Figure 4A) to farther investigate the origin of this specific intensity profile and compare it to membrane proteins. These selected soluble proteins have fewer alpha helices than other soluble proteins, and have more beta strands and turns (Figure 4B,C). This observation reaches the conclusion drawn from Figure 3B, that soluble proteins with higher content in beta strands show a peak in intensity profile. Separation of the intensity profile by percentage of secondary structure (Figure 4D,E,F) shows that the percentage of secondary structure doesn’t influence the peak hight, and is more visible on the inflection around 6.5Å resolution. The behavior of these selected soluble proteins thus does not exactly match the one of membrane proteins, leading to think that the reasons for the appearance of this peak originate from different sources.

**Figure 4.**
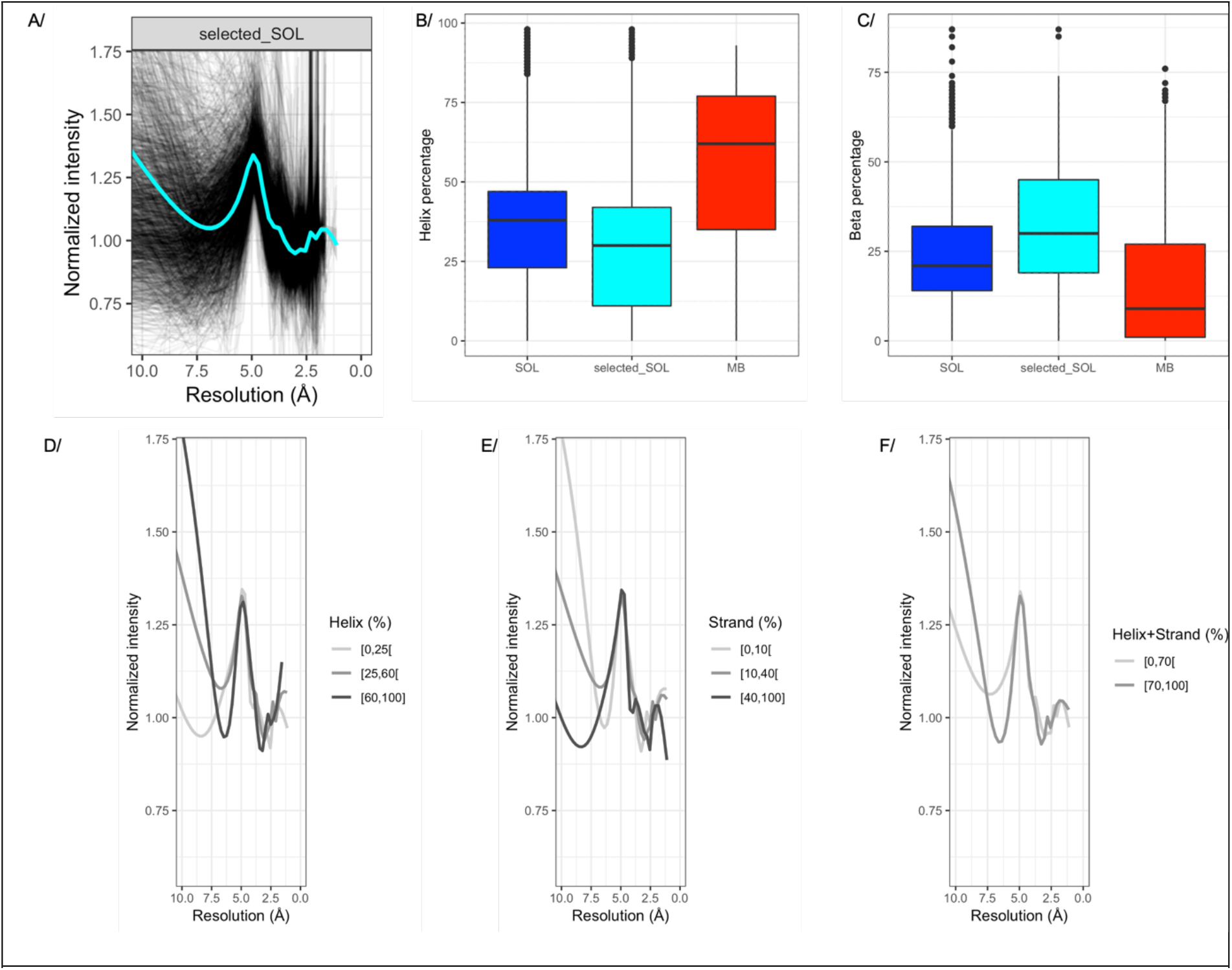
Soluble proteins showing a peak in intensity profile and relation to secondary structure elements. A/ Intensity profile and smoothed profile (cyan) for soluble proteins selected to have an intensity profile greater than 1.25 at 4.9Å resolution. Proportion of alpha helices (B/) or Beta strands (C/) in soluble (SOL) and selected soluble (selected_SOL) proteins *vs* membrane proteins (MB). D/E/F/ overlaid smoothed intensity profiles separated by alpha helices percentage, beta strands percentage of the sum of secondary structures.

### 3.5. The peak centred on 4.9 Å resolution originates from alpha helices and beta sheets collinearity in membrane proteins, and high content beta sheets for selected soluble proteins

The fact that embedded membrane proteins display a very marked peak in their intensity profiles, as shown in Figure 2, is remarkable. Indeed, these proteins have been selected to be only present in the membrane (Robert *et al*., 2018, 2017), with no outside domains protruding in the periplasm or cytoplasm. They are constituted of alpha helices or beta strands oriented perpendicular to the membrane plane. This observation led us to investigate the influence of secondary structure collinearity on intensity profiles. DSSP parameters (Touw *et al*., 2015) were extracted for each entry, and collinearity index of secondary structures calculated. A collinearity value of 0.3 for alpha helices or beta strands means a random orientation of these secondary structure elements. Membrane proteins have very collinear alpha helices with most of the values above 0.6 (Figure 5A). There is also a cluster at high helix proportion and high collinearity, meaning that when there is a high proportion of alpha helices, they tend to distribute in a colinear fashion. This trend is confirmed for embedded membrane proteins. Soluble proteins have much lower collinearity, with most of the values distributing below 0.6, with a well-formed cluster around 40% helices and 0.5 collinearity index, denoting a similar distribution of alpha helices across all the soluble protein structures. Beta strands for membrane proteins show two peaks, centred on 25 and 55%. The first peak corresponds to the distribution seen for soluble proteins (Figure 5B). The second peak for higher content of beta sheets in membrane proteins corresponds to proteins fully embedded in the membrane, folding as beta barrels(Robert *et al*., 2017). The strand collinearity is thus a little greater, being normal to the membrane plane. For selected soluble proteins, showing a greater content of beta strands, they have a bimodal distribution of beta strands resembling the one of membrane proteins in terms of secondary structure elements proportion (Figure 5B). The strand collinearity index is however similar in both distributions. We confirmed these observations by exploring the influence of secondary structure collinearity on the height of the peak at 4.9 Å resolution (Figure 5D,E). For membrane proteins, there is a clear trend to a higher peak with more collinearity of alpha helices or beta strands. For soluble proteins, collinearity has no influence on the peak hight.

**Figure 5.**
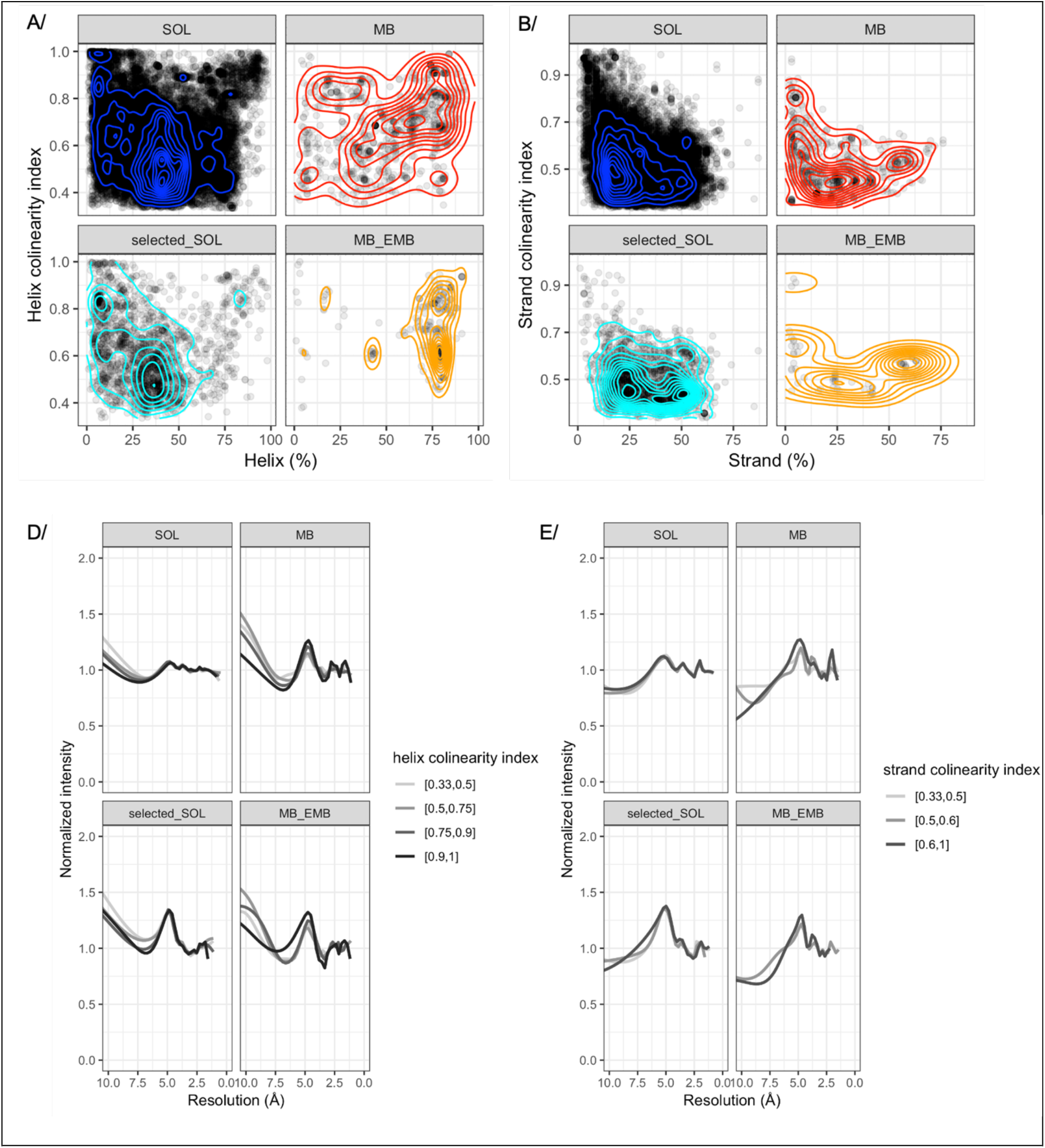
Collinearity of secondary structures. A/ distribution of alpha helices collinearity as a function of helix proportion. Each entry is represented by a transparent circle for clarity. Level lines help to identify clusters and more populated zones. The panel is divided and color coded as the other figures above. B/ same as A/ on strand collinearity. D/ E/ influence of helix and strand collinearity respectively on diffraction intensity profiles.

The reasons for the peak in intensity profiles is thus different for both classes of proteins. Membrane proteins have more alpha helices than soluble ones, and these helices are more collinear. When the peak is due to beta sheets, it is also because these beta sheets are oriented. Soluble proteins with a peak in intensity profiles have larger proportion of beta sheets, not necessarily more collinear than soluble protein without the peak, but this larger proportion results in a different intensity profile.

### 3.6. Proteins showing the peak at 4.9Å resolution correspond to a higher diffraction anisotropy population

Finally, the relationship between the presence of a peak in intensity profile and diffraction anisotropy was explored. Diffraction anisotropy was calculated for each entry and the population represented in density as a function of diffraction anisotropy. Each class of protein was separated for analysis. Importantly, since the relationship between diffraction anisotropy and resolution of the dataset has been clearly described(Robert *et al*., 2017), the current data was separated by resolution range (Figure 6). Interestingly, selected soluble proteins follow the same distribution as membrane proteins, corresponding to a higher diffraction anisotropy distribution compared to soluble proteins. This is a first observation of a physical characteristic that can isolate a population of soluble proteins that resemble membrane proteins in terms of anisotropy distribution. This observation also links diffraction anisotropy to the peak in intensity profiles. Of course, this link should also be put in perspective of the anisotropy distribution that shows a large scattering of points with a fair number of structures with low anisotropy.

**Figure 6.**
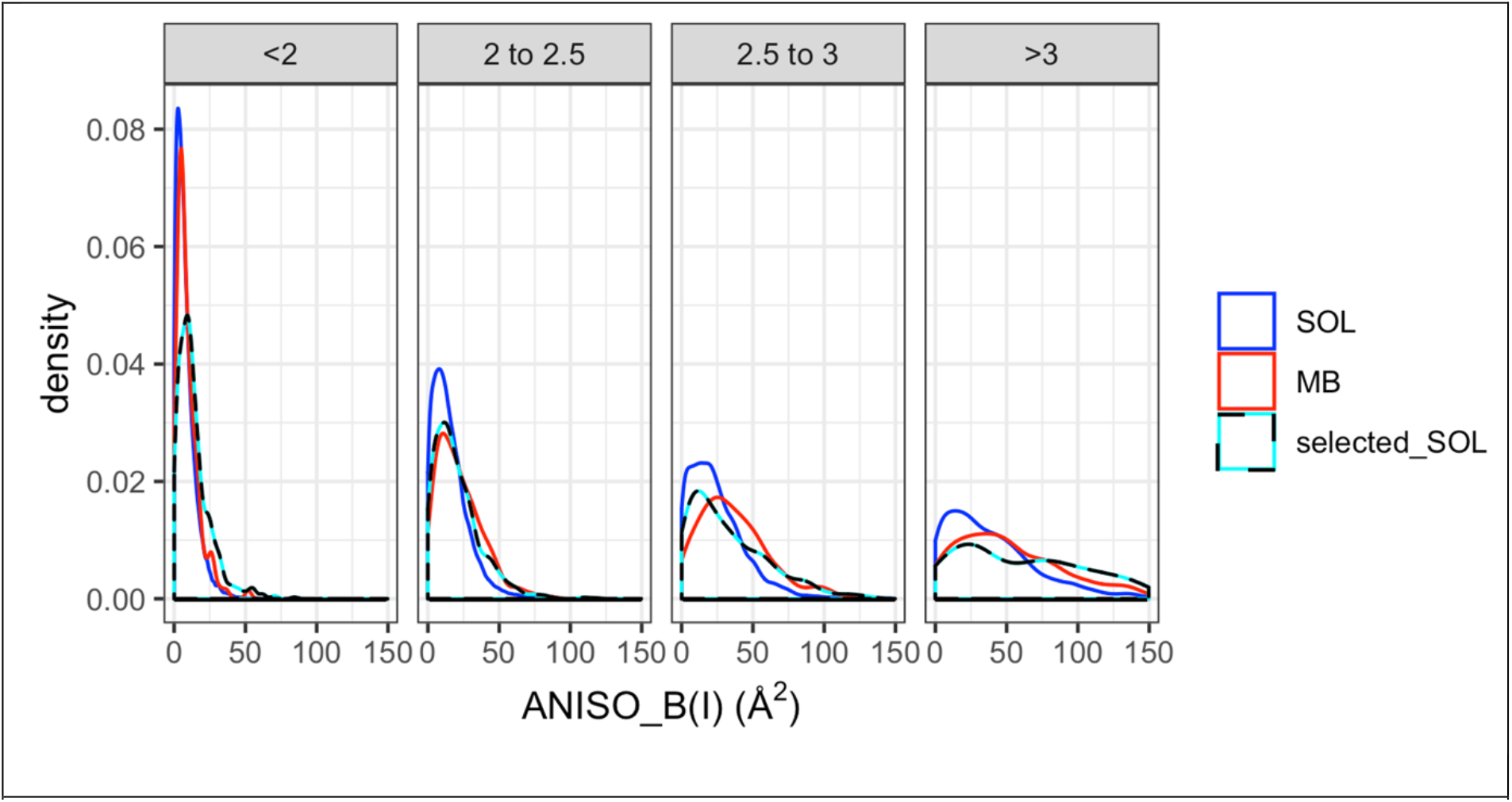
Link between diffraction anisotropy and presence of a peak at 4.9 Å in intensity profiles. Distribution of anisotropy values calculated on intensities (ANISO_B(I)) are separated by resolution ranges: less than 2Å, between 2 and 2.5Å, between 2.5 and 3Å and greater than 3Å resolution. Distributions for soluble proteins are in blue, for membrane proteins in red and for selected soluble proteins in dotted cyan-black lines. Embedded membrane proteins were omitted due to the low number of structures in the current database.

### 3.7. Discussion

Diffraction anisotropy is a very prevalent problem in membrane protein crystallography, but the root of this problem still remains unknown, and the reason why membrane proteins display more anisotropy than soluble proteins is still in the dark.

Here we show that diffraction anisotropy and difference in diffraction limits are two separate, yet coupled, issues. They often tend to appear in tandem, which is the origin of the claims that the difference in diffracting limits causes diffraction anisotropy(Strong *et al*., 2006, Dauter *et al*., 2005, Sheriff & Hendrickson, 1987). At the core, difference in diffraction limits comes from the absence of crystal contacts in specific directions of space, resulting in more order and thus stronger diffraction(Sheriff & Hendrickson, 1987, Chaptal *et al*., 2016). Diffraction anisotropy itself originates from a deviation from an experimental curve obtained from a set of crystals diffracting at high resolution, the BEST curve(Popov & Bourenkov, 2003). Often these two phenomena appear in concert, but not always as judged by the large scattering of data points in Figure 1, and the large variation of anisotropy seen when there is no difference in resolution limits. There is nevertheless a trend to more anisotropy with more difference in resolution limits, coupling the two otherwise-separated events. On top of it, these two events occur more prevalently on low resolution datasets, which increases the effects and lowers electron density map qualities. The lack of completeness in the highest resolution shells due to difference in resolution limits increases noise in maps, and diffraction anisotropy creates artifacts(Global_Phasing_Limited, 2018).

It has been clearly established that membrane proteins harbour more diffraction anisotropy, regardless of their crystallization techniques (in detergent or *in-meso*), of their insertion in the membrane, of their function, nor if they are fully embedded in the membrane or having large parts protruding outside(Robert *et al*., 2017). It is their membranous nature that give membrane proteins their capability to diffract X-rays more anisotropically compared to soluble proteins. The insertion into the membrane is made exclusively by alpha helices or by beta strands forming a beta barrel. Secondary structure elements are thus are the centre of attention. We analysed diffraction intensity profiles and identified a peak centred on 4.9 Å resolution, a zone previously annotated to correspond to secondary structure elements. We showed that the proportion of secondary structure elements is not enough to attest for the presence of this peak, and of its height. Following the idea brought by fully embedded membrane proteins that show a very directional fold, normal to the membrane plane, we explored the influence of secondary structure collinearity on the presence of the peak. We could show that membrane protein’s secondary structures distribute more collinearly, and that the peak increases in hight with increasing secondary structures collinearity; the more collinear alpha helices or beta sheets (in a barrel), the higher the peak at 4.9 Å resolution (Figure 5). For soluble proteins however, such trend could not be seen, likely due to the fact that secondary structure elements tend to distribute more “randomly” than for membrane proteins.

We could then establish a clear link between the peak at 4.9 Å in intensity profile and a protein distribution to higher anisotropy (Figure 6). Selected soluble proteins for a high peak in intensity profile follow the same distribution as membrane proteins, that always have the peak. This observation is definitely intriguing, allowing for the first time to extrapolate a set of soluble proteins following the anisotropic distribution of membrane proteins. Acknowledgedly, the present analysis was performed in one dimension, averaging the 3D intensity profiles, and should be continued in 3D by scrutinizing intensity profiles following the anisotropy ellipsoids(Global_Phasing_Limited, 2018). Since diffraction anisotropy is occurring in 3D, being different on every cell axis or directions of the diffraction ellipsoid, it would be interesting to investigate the intensity profiles of these different classes of proteins as a function of the ellipsoid main axis, in a follow-up article.

An additional important consideration is that an X-ray diffraction dataset is obtained from a rotation of the crystal in respect to a fixed axis linking the X-ray beam to the detector. We showed here that secondary structure collinearity is an important factor linked to anisotropy, but there are also other factors at play as shown by the distribution of anisotropy (Figure 6). The root of diffraction is scattering of individual elements, repeated in space and thus amplified by the spatial arrangement in the crystal. It is tempting to speculate that the source of diffraction anisotropy is collinearity of secondary structure, not only within the protein, but collinearity of secondary structures in the whole crystal in respect to the X-ray beam-detector axis. These orientations are found more prevalently in membrane proteins for two reasons: 1/ membrane proteins have intrinsically more collinear structures and 2/ their hydrophobic belts prevents crystal contacts, thus orienting crystal contacts in some places leading to more overall collinearity. All together, these data bring to light the fact that the special 3D arrangement of membrane proteins brought by the membrane insertion induces a specific X-ray diffraction. Just like their membrane insertion calls for specific tools to purify membrane proteins (detergents, amphipols, polymers, nanodiscs, etc…), the fact that they behave differently than soluble proteins in regard to biophysical experiments calls for a specific data handling for membrane proteins in general.

## Supporting information

Raw data used in the analysis

## Acknowledgements

The authors wish to thank Randy Read for pointing to the ANO mode of Phaser so that the intensity profiles can be exploited and the present data analysed, and for helpful discussions in preparing this manuscript.

## Funding

This work was supported by the *Centre National de la Recherche Scientifique* (CNRS) and Lyon University for salaries. The authors also want to acknowledge ANR-19-CE11-0023-01.

## Notes

### Competing Interest Statement

The authors have declared no competing interest.

## References

Bayburt, T. H. & Sligar, S. G. (2010). FEBS Lett 584, 1721–1727.

Berman, H. M., Westbrook, J., Feng, Z., Gilliland, G., Bhat, T. N., Weissig, H., Shindyalov, I. N. & Bourne, P. E. (2000). Nucleic Acids Research 28, 235–242.

Caffrey, M. (2015). Acta crystallographica. Section F, Structural biology communications 71, 3–18.

Chacko, A. R., Zwart, P. H., Read, R. J., Dodson, E. J., Rao, C. D. & Suguna, K. (2012). Acta Crystallogr D Biol Crystallogr 68, 1541–1548.

Chae, P. S., Rasmussen, S. G., Rana, R. R., Gotfryd, K., Chandra, R., Goren, M. A., Kruse, A. C., Nurva, S., Loland, C. J., Pierre, Y., Drew, D., Popot, J. L., Picot, D., Fox, B. G., Guan, L., Gether, U., Byrne, B., Kobilka, B. & Gellman, S. H. (2010). Nat Methods 7, 1003–1008.

Chaptal, V., Kilburg, A., Flot, D., Wiseman, B., Aghajari, N., Jault, J. M. & Falson, P. (2016). Data Brief 7, 726–729.

Dalton, J. A., Michalopoulos, I. & Westhead, D. R. (2003). Bioinformatics 19, 1298–1299.

Dauter, Z., Botos, I., LaRonde-LeBlanc, N. & Wlodawer, A. (2005). Acta Crystallogr D Biol Crystallogr 61, 967–975.

Dörr, J. M., Scheidelaar, S., Koorengevel, M. C., Dominguez, J. J., Schäfer, M., van Walree, C. A. & Killian, J. A. (2016). Eur Biophys J 45, 3–21.

Dufourc, E. J. (2021). Biochimica et biophysica acta. Biomembranes 1863, 183478.

Evans, P. R. & Murshudov, G. N. (2013). Acta Crystallogr D Biol Crystallogr 69, 1204–1214.

Global_Phasing_Limited (2018). The Staraniso Server, http://staraniso.globalphasing.org/cgi-bin/staraniso.cgi.

Johansen, N. T., Luchini, A., Tidemand, F. G., Orioli, S., Martel, A., Porcar, L., Arleth, L. & Pedersen, M. C. (2021). Langmuir.

Kabsch, W. & Sander, C. (1983). Biopolymers 22, 2577–2637.

Kozma, D., Simon, I. & Tusnády, G. E. (2013). Nucleic Acids Res 41, D524–529.

Krishnarjuna, B., Ravula, T. & Ramamoorthy, A. (2020). Chemical Communications 56, 6511–6514.

Marconnet, A., Michon, B., Le Bon, C., Giusti, F., Tribet, C. & Zoonens, M. (2020). Biomacromolecules.

Matar-Merheb, R., Rhimi, M., Leydier, A., Huche, F., Galian, C., Desuzinges-Mandon, E., Ficheux, D., Flot, D., Aghajari, N., Kahn, R., Di Pietro, A., Jault, J. M., Coleman, A. W. & Falson, P. (2011). PLoS One 6, e18036.

McCoy, A. J., Grosse-Kunstleve, R. W., Adams, P. D., Winn, M. D., Storoni, L. C. & Read, R. J. (2007). J Appl Crystallogr 40, 658–674.

Morris, R. J., Blanc, E. & Bricogne, G. (2004). Acta Crystallogr D Biol Crystallogr 60, 227–240.

Morris, R. J. & Bricogne, G. (2003). Acta Crystallogr D Biol Crystallogr 59, 615–617.

Nguyen, K. A., Peuchmaur, M., Magnard, S., Haudecoeur, R., Boyere, C., Mounien, S., Benammar, I., Zampieri, V., Igonet, S., Chaptal, V., Jawhari, A., Boumendjel, A. & Falson, P. (2018). Angew Chem Int Ed Engl 57, 2948–2952.

Oluwole, A. O., Danielczak, B., Meister, A., Babalola, J. O., Vargas, C. & Keller, S. (2017). Angew Chem Int Ed Engl 56, 1919–1924.

Overduin, M. & Esmaili, M. (2019). Chemistry and Physics of Lipids 218, 73–84.

Popov, A. N. & Bourenkov, G. P. (2003). Acta Crystallogr D Biol Crystallogr 59, 1145–1153.

R Core Team (2016). R: A language and environment for statistical computing.

Ravula, T., Hardin, N. Z. & Ramamoorthy, A. (2019). Chemistry and Physics of Lipids 219, 45–49.

Read, R. J. & McCoy, A. J. (2016). Acta crystallographica. Section D, Structural biology 72, 375–387.

Robert, X., Kassis-Sahyoun, J., Ceres, N., Martin, J., Sawaya, M. R., Read, R. J., Gouet, P., Falson, P. & Chaptal, V. (2017). Scientific reports 7, 17013.

Robert, X., Kassis-Sahyoun, J., Ceres, N., Martin, J., Sawaya, M. R., Read, R. J., Gouet, P., Falson, P. & Chaptal, V. (2018). Data Brief 19, 753–757.

Russo Krauss, I., Merlino, A., Vergara, A. & Sica, F. (2013). Int J Mol Sci 14, 11643–11691.

Salnikov, E. S., Aisenbrey, C., Anantharamaiah, G. M. & Bechinger, B. (2019). Chemistry and Physics of Lipids 219, 58–71.

Sawaya, M. R. (2014). Methods Mol Biol 1091, 205–214.

Sheriff, S. & Hendrickson, W. A. (1987). Acta Crystallogr A 43, 118–121.

Strong, M., Sawaya, M. R., Wang, S., Phillips, M., Cascio, D. & Eisenberg, D. (2006). Proc Natl Acad Sci U S A 103, 8060–8065.

Touw, W. G., Baakman, C., Black, J., te Beek, T. A., Krieger, E., Joosten, R. P. & Vriend, G. (2015). Nucleic Acids Res 43, D364–368.

Wickham, H. (2016). Springer-Verlag New York.

Winn, M. D., Ballard, C. C., Cowtan, K. D., Dodson, E. J., Emsley, P., Evans, P. R., Keegan, R. M., Krissinel, E. B., Leslie, A. G., McCoy, A., McNicholas, S. J., Murshudov, G. N., Pannu, N. S., Potterton, E. A., Powell, H. R., Read, R. J. Vagin, & Wilson, K. S. (2011). Acta Crystallogr D Biol Crystallogr 67, 235–242.

wwPDB_consortium (2018). Nucleic Acids Res.

Zwart, P. H. & Lamzin, V. S. (2003). Acta Crystallogr D Biol Crystallogr 59, 2104–2113.

